# Prospective and retrospective contributions to intention awareness in voluntary action

**DOI:** 10.1101/821488

**Authors:** Matthias Schultze-Kraft, Elisabeth Parés-Pujolràs, Karla Matić, Patrick Haggard, John-Dylan Haynes

## Abstract

Spontaneous, voluntary actions are often accompanied by motor intentions, that is the experience of “being about to move”. The emergence of this experience is contentiously discussed, culminating in two opposing views of motor intention. While prospective theorists hold that an intention to move genuinely precedes the action, retrospective theorists suggest that an intention is reconstructed only after action execution. As such, both theories offer radically different predictions about the possibility of voluntary control over action intention. We assess here the respective contributions of prospective and retrospective factors in motor intention awareness in a real-time EEG setup. Participants performed self-paced movements with intermittent interruption by cues, signalling to either move or not to move. Cues were triggered by a brain-computer interface upon detection of either the presence or absence of a readiness potential (RP), a signal known to precede self-paced voluntary actions. Subsequently, participants reported their intention to move at the time of the cue. We found that participants were more likely to report an intention to move when the cue was followed by a movement than no movement, highlighting the effect of retrospection on awareness of intention. Interestingly, however, an intention to move was also more frequently reported when the cue was elicited by an RP, compared to when no RP was present, suggesting that awareness of intention is also informed by motor preparation processes, congruent with prospective theories. Overall, our findings show that intention awareness in voluntary action relies on both prospective and retrospective components, and we describe here how these two factors are dynamically integrated over time.

## 1. Introduction

When we perform spontaneous, voluntary movements, our subjective experience is that of a coherent flow of conscious events, from forming the intention to act to executing the movement. In a similar vein, neurophysiological data suggest a succession of neural events, with the execution of voluntary movements being preceded by brain signals that indicate motor preparation (Kornhuber and Deecke, 1965; Shibasaki and Hallett 2006). Despite extensive research on the relationship between awareness of intention and motor preparatory processes, it remains unclear when an intention to move becomes accessible to consciousness. One line of research suggests that we have conscious access to our motor preparation processes and thus become aware of our intention to move *before* action initiation (Libet et al. 1983; Haggard and Eimer 1999; Matsuhashi and Hallett 2008; Parés-Pujolràs et al. 2019). That is, we consciously think about what we are about to do before actually doing it (*prospective hypothesis*). In contrast, others have argued that intentions are postdictively inferred *after* action initiation (Dennett and Kinsbourne, 1992; Wegner, 2002; Kühn and Brass 2009). In this view, it is the fact of having done something that generates the idea that we intended to do it (*retrospective hypothesis*). An intriguing third possibility is an integrative model that views awareness of intention as a process extended in time during which prospective and retrospective effects might be integrated (Lau et al. 2007; Douglas et al. 2015; Verbaarschot et al. 2016). Comparator models of action control suggest that efferent copies of motor commands and sensory feedback after action execution are integrated, and these efferent copies are postulated to carry different types of information about the movement characteristics, including *when* it will be executed (Wolpert and Kawato 1998). We suggest that a similar integration mechanism is in play for intention awareness that is over and above the prospective and retrospective effects. We refer to this as the *temporal integration hypothesis*.

In this study, we investigate how motor preparation processes and action execution interact over time to produce the experience of intention that accompanies spontaneous movements. We approached this by combining real-time EEG-monitoring of a self-paced task with a classical *Go*/*No-Go* task. We aimed to interrupt participants with Go/No-Go cues either while they were *preparing* to execute a self-paced movement, or at a time when they were not preparing at all. After each interruption, we asked them about their intention to move. Our real-time method allowed us to investigate in an unbiased way how motor preparatory states influence conscious intentions.

One candidate motor preparation signal is the so-called readiness potential (RP), an increasing negativity over the motor cortex that was first shown to precede self-paced movements by Kornhuber and Deecke (1965). The RP has later been linked to awareness of intention (Libet et al. 1983; Haggard and Eimer 1999; Sirigu et al. 2004; Schlegel et al. 2013; Frith and Haggard 2018; Parés-Pujolràs et al. 2019). While the RP can be observed as early as 2s before movement execution, people generally report becoming aware of an impending movement only about 200 ms before movement onset (Libet et al., 1983). Although concerns have been raised about the validity of the RP as a neural signal uniquely linked to voluntary motor preparation (Schurger et al. 2012; Jo et al. 2013), recent evidence shows that it can be used to predict voluntary movement at a single trial level (Schultze-Kraft et al. 2016). This does not rule out the possibility that RP-like activity can be found in the absence of an action, but does suggest that the presence of an RP reflects an increased probability of action. The RP remains one of the best candidate signals providing information about motor preparation, and is thus a potential prospective cause of intention awareness.

In the current study, participants performed a self-paced task during which they were instructed to press a footpedal at any time they wished after trial onset. Occasionally, they would be interrupted by either a green (*Go*) or a red (*No*-*Go*) cue, instructing them to press the pedal immediately or inhibit any movement, respectively. Importantly, cues were triggered either at a random time when no RP was detected (*RP*−), or as soon as a readiness potential was detected (*RP*+) by a brain-computer interface (BCI) monitoring participants’ electroencephalogram (EEG) in real-time. Each trial was randomly assigned a combination of motor preparation state (*RP*+/*RP*−) and action execution instruction (*Go*/*No*-*Go*). When interrupted by a cue, participants were given time to respond accordingly (execute / inhibit a movement) and were additionally asked to verbally report (*Yes/No*) whether they were preparing to move at the time the coloured cue was presented.

Our experimental design allowed us to directly test several predictions that follow from the hypotheses formulated above. First, if people have conscious access to motor preparatory processes, awareness of intention should be more likely in the presence of a signal indicating motor preparation. Thus, the prospective hypothesis predicts a higher rate of awareness reports when probed in the presence of an RP than when probed in the absence of an RP. Second, the retrospective hypothesis states that it is the action execution itself that produces the experience of intention. Hence, it predicts that participants should be more likely to report a prior intention if an action is eventually executed (i.e. reports following a green *Go* cue) than when it is not (i.e. reports following a red, *No*-*Go* cue). Since the colour of the cue is independent of the presence/absence of an RP, these two predictions are independent. Finally, the temporal integration hypothesis proposes that intention judgements depend on the integration of motor representations preceding action and on sensory feedback after action execution. We predict that prospective information about motor preparation based on transient signals such as the RP is only available for integration with (retrospective) sensory feedback for a short time. Thus, we expect that the effect of motor preparation processes on intention judgements should be stronger if a movement is executed shortly after the presence of an RP.

## 2. Methods

### 2.1. Participants

We investigated 23 healthy, naive participants (17 female, mean age 30, SD 5.2 years). The experiment was approved by the local ethics board and was conducted in accordance with the Declaration of Helsinki. All participants gave their informed oral and written consent.

### 2.2. Experimental setup

Participants were seated in a chair facing a computer screen at a distance of approximately 1m. They were asked to place their hands in their lap and their right foot 2 cm to the right of a 10 cm × 20 cm floor-mounted switch pedal (Marquardt Mechatronik GmbH, Rietheim-Weilheim, Germany). Throughout the experiment, EEG was recorded at 1 kHz with a 64-electrode Ag/AgCl cap (EasyCap, Brain Products GmbH, Gilching, Germany) mounted according to the 10-20 system and referenced to FCz and re-referenced offline to a common reference. Given that the signal of interest, the readiness potential, is observed predominantly over central electrodes, EEG was recorded from the following 30 electrodes: F3, F1, Fz, F2, F4, FC5, FC3, FC1, FC2, FC4, FC6, C5, C3, C1, Cz, C2, C4, C6, CP5, CP3, CP1, CPz, CP2, CP4, CP6, P3, P1, Pz, P2, P4. In addition to the EEG, the right calf electromyogram (EMG) was recorded using surface Ag/AgCl electrodes in order to obtain the earliest measure of movement onset. The amplified signal (analog filters: 0.1, 250 Hz) was converted to digital (BrainAmp MR Plus and BrainAmp ExG, Brain Products GmbH, Gilching, Germany), saved for offline analysis, and simultaneously processed online by the Berlin Brain-Computer Interface Toolbox (BBCI, github.com/bbci/bbci_public). The Pythonic Feedback Framework (Venthur et al. 2010) was used to generate visual feedback. Verbal reports in response to the prompting task (see below) were recorded by a microphone that was placed on the table and manually transcribed trial-by-trial after the experiment. Verbal reports were chosen over movement reports to disentangle the motor signal effects used in the main motor task (see below) from the intention reports.

### 2.3. Experimental design and task

The experiment was divided in three stages (Fig 1). In a preparatory experimental stage I, participants performed a simple self-paced task. The data collected in stage I were used to train a classifier to monitor EEG activity in real-time during stage II. In stage II, the main experiment, participants performed the same self-paced task and a prompting task. In a supplemental stage III, participants performed a cued reaction task.

**Figure 1:**
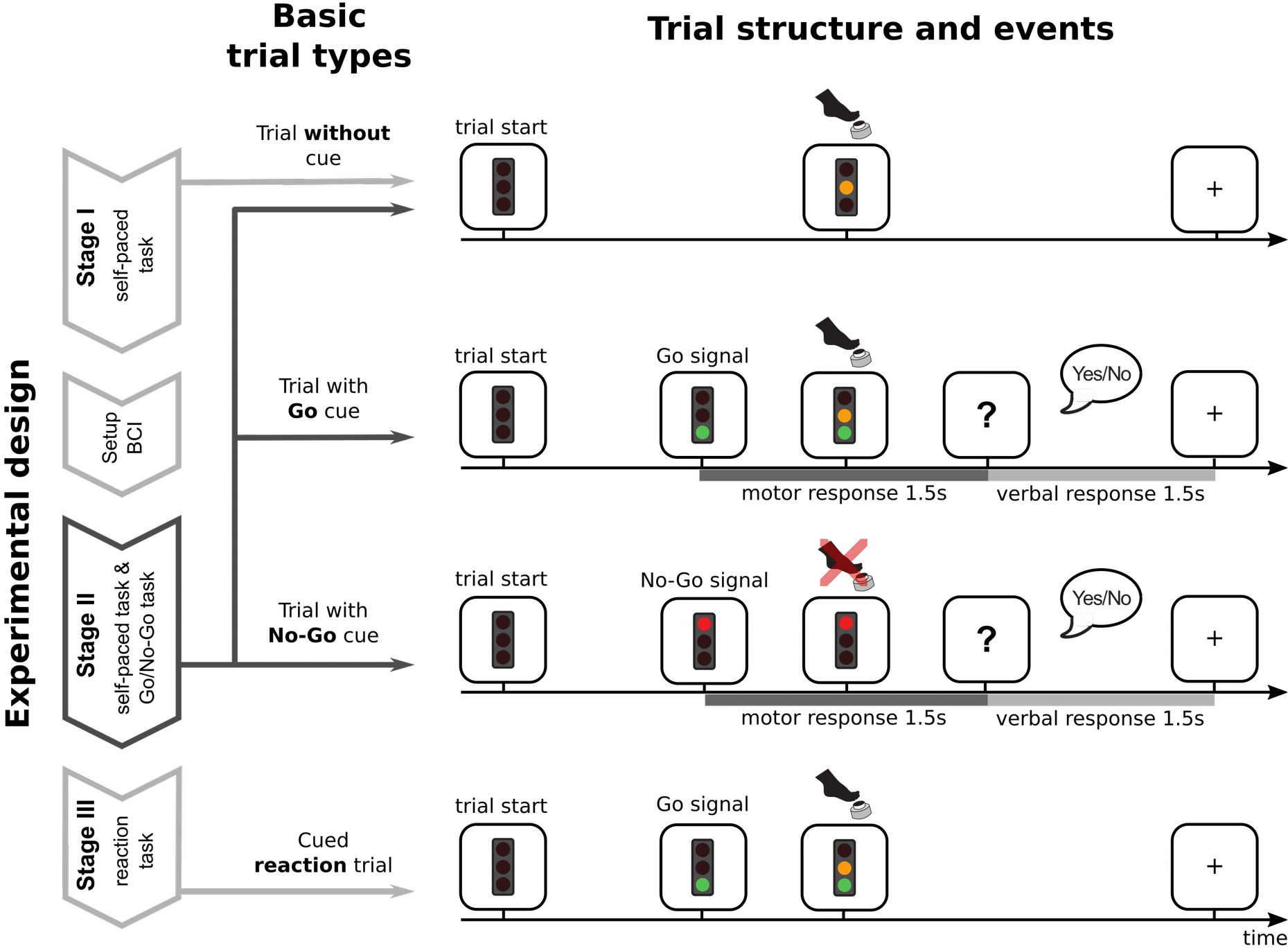
Experimental design and trial types and events. During stage I, participants performed spontaneous, self-paced pedal presses in single trials. No green or red cues were elicited and a trial always ended with a pedal press. After stage I, a BCI was trained on EEG data recorded during stage I. During stage II, the main experiment, participants again performed spontaneous, self-paced pedal presses, but were occasionally interrupted by either the green or the red light turning on. If they pressed the pedal before either light was turned on, the trial ended as in stage I. If the green or the red light was turned on, participants had 1.5 seconds to follow the instruction, i.e. to press immediately after a green cue or not press / inhibit after a red cue. Subsequently, the question “Were you about to press?” appeared on screen for 1.5 seconds (indicated by a question mark) during which participants were asked to verbally respond. In stage III, participants performed a simple reaction task: In each trial, the green light would turn on after a random time (chosen from a uniform distribution between 2 and 5 seconds), after which participants were instructed to press the pedal as quickly as possible.

#### 2.3.1. Stage I: Collection of training data for the classifier

During stage I, participants performed a simple self-paced task. The start of a trial was signaled by a traffic light display appearing on the screen with all three colored lights (green, yellow, red) turned off. Participants were instructed to wait for roughly 2 seconds, after which they could press the pedal at any time. They were asked to avoid preplanning the movement, avoid any obvious rhythm, and to press when they felt the spontaneous urge to move (Kornhuber and Deecke, 1965; Libet et al., 1983). When the pedal was pressed the yellow light was turned on for 1 second, after which the traffic light disappeared and was replaced by a fixation cross. This constituted the end of a trial. The fixation cross remained onscreen for a 3s intertrial period. Participants were asked to fixate and remain relaxed without moving. Each participant performed a total of 100 trials in stage I, with the possibility of taking a break after each 25 trials.

#### 2.3.2. Stage II: Main experiment

In stage II, participants performed the same self-paced task indicated above, but additionally they would sometimes be interrupted by either the green (*Go*) or red (*No*-*Go*) traffic light turning on for a duration of 1.5 s before the trial ended. Participants were instructed to press the pedal as quickly as possible in response to the green light, and to withhold from moving or to abort any potentially planned pedal press in response to the red light. They were given 1.5 s to respond to this Go/No-Go task. A *Go* trial was considered correct if the pedal was pressed while the green light was on. When participants pressed the pedal, the yellow traffic light turned on for 1 second. A *No*-*Go* trial was considered correct if the pedal was not pressed while the red light was on. If a trial was not executed correctly, it ended with a fixation cross and was discarded from further analysis. After correct trials, the question “*Were you about to press?”* appeared on screen for 1.5 s. Participants were instructed to verbally report (*”Yes”*/*”No”*) whether they were preparing to move at the time the coloured cue appeared on screen.

Each trial in stage II was randomly assigned to one of four conditions defined by a combination of two factors. The first factor was the Action execution instruction (*Go*/*No*-*Go*), while the second one was the motor preparation state that would be used to trigger the instruction (*RP*+/*RP*−). Thus, while the former determined *which* light would be turned on in the trial (green in Go trials and red in *No*-*Go* trials), the motor preparation state determined *when* the light would be turned on. Note that this assignment of trials was putative rather than absolute, because participants sometimes performed self-paced movements, as in stage I, before they were interrupted by any cue. This was a consequence of how the timing of the cues was realized in *RP*+ and *RP*− trials, which we describe in details in section ***2.4***. Stage II had a total duration of 60 minutes, with the possibility to take a break every 15 minutes. The number of trials executed in stage II varied across subjects depending on the frequency of participants’ self-paced actions and the time at which the cues were presented.

#### 2.3.3. Stage III: Supplementary task

In stage III, participants performed a simple, cued reaction task. The start of a trial was signalled by the traffic light display appearing on the screen with all three coloured lights turned off. In each trial, the green light would turn on for 1.5 seconds after a random time, chosen from a uniform distribution between 2 and 5 seconds. Participants were instructed to respond as fast as possible with a pedal press as soon as the green light appeared. When they pressed the pedal, the yellow light was turned on for 1 second after which the traffic light disappeared and was replaced by a fixation cross. This constituted the end of a trial. The fixation cross remained onscreen for a 3s intertrial period. Each participant performed this task for a total time of 8 minutes, with the possibility to take a break after 4 minutes. The aim of this stage was to obtain measures of speeded reaction times in the absence of a self-paced task, and to compare them to the reaction times to *Go* cues obtained in stage II.

### 2.4. Real-time BCI predictor

For the BCI predictor used in stage II, a linear classifier was trained on EEG data from the 100 pedal presses recorded during stage I.

#### 2.4.1. EMG onset detection

For each trial, we assessed the movement onset. For higher temporal precision, we defined the onset of the movement based on the EMG rather than based on the final completion of the movement with the pedal press. To obtain EMG onset we high-pass filtered the EMG signal at 20 Hz. Then the standard deviation of the signal during the first 1000 ms after each trial start cue was determined as an “idle” baseline. For each trial individually, the standard deviation of subsequent, overlapping 50 ms windows was computed and EMG onset identified as the end of the first 50 ms window where the standard deviation exceeded idle baseline by a factor of 3.5.

#### 2.4.2. Class specification

Based on these movement onsets, two periods were defined as “Move” and “Idle” for the training of the classifier. The *Move* periods were 1200 ms long segments preceding EMG onset, while the *Idle* periods were 1200 ms long segments preceding the trial start cue (i.e. the onset of the traffic light).

#### 2.4.3. Feature extraction and classifier training (Stage I)

EEG data from those segments were baseline corrected to the mean signal in the time interval between −50 and 0 ms w.r.t. EMG onset or trial start cue, respectively. These were then averaged over time windows defined by the time points −1200, −900, −650, −450, −300, −200, −100 and −50 ms w.r.t. EMG onset or trial start cue, respectively. The choice of the baseline correction interval being locked to the end of the EEG segment (as opposed to the traditional choice of being locked to the beginning of the segment) and the choice of unequal time intervals were both based on a piloting analysis on previous data (Schultze-Kraft et al., 2016) that showed improved classification accuracy with these parameters. The resulting values were concatenated and used as features to train a regularized Linear Discriminant Analysis (LDA) classifier with automatic shrinkage (Blankertz et al. 2011). Classification accuracy (obtained from a 10-fold cross-validation on stage I data) was > 70% for all participants.

#### 2.4.4. Real-time application of classifier (Stage II)

During stage II, the so-trained classifier was used to monitor the ongoing EEG in real-time. Therefore, every 10 ms a feature vector was constructed from the immediately preceding 1200 ms of EEG data, as outlined above, and used as input to the classifier, generating a classifier output value every 10 ms. This output variable was a continuous signal that probabilistically classified the current EEG segment either to the *Idle* or to the *Move* class.

#### 2.4.5. Threshold setting

A classifier threshold was set for each participant individually. Because the classifier output signal was likely to mirror the stochastic nature of the EEG, a conservative threshold was defined in order to avoid many cues to be prematurely triggered by noise. For this, we trained the classifier on 99 trials from stage I and applied it to each consecutive and overlapping 1200 ms feature window in the left out trial, thereby mimicking the real-time application during stage II. This was for done for each of the 100 trials in a leave-one-out crossvalidation scheme. For each of these continuous classifier output vectors the time of first threshold crossing after trial start was computed. Let us refer to the time of first threshold crossing in a trial as a “prediction” event. Now, we define predictions occurring somewhere between trials start and up to 600 ms before movement onset as false alarms (FA), predictions occurring between 600 ms before movement onset and the time of movement onset as Hits, and predictions occurring after movement onset or not occurring at all as Misses. From this the F-measure (Powers 2011) 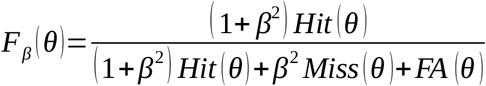 was computed for different threshold values θ. The largest F thus corresponds to the threshold θ were the Hit rate is maximal, while at the same time the FA and Miss rates are minimal. Moreover, by choosing *β*=0.5, we aimed at giving the minimization of FAs more weight than minimizing Miss rate. We prioritized minimizing the number of false alarms, at the cost of potentially missing some actions. The resulting F values from stage I were smoothed and the threshold with the highest F value chosen.

### 2.5. Timing of cues during stage II

#### 2.5.1. Timing of *RP*+ cues

During stage II, if a trial was assigned as an *RP*+ trial, the BCI was inactive during the first 1500 ms after trial start. This ensured that an RP+ cue was not elicited during the minimum self-paced waiting time of 2 sec instructed to participants. After 1500 ms, the BCI was activated and either the green or the red light were turned on as soon as the classifier reached the specific threshold, indicating that the current EEG data had changed from class “Idle” to class “Move”.

#### 2.5.2. Timing of *RP*− cues

During stage II, if a trial was assigned as a *RP*− trial, a cue was elicited after a predefined random time that was chosen before trial start for each trial individually. In these trials, to ensure that the cue was displayed at a plausible time given behavioural characteristics of the participant, a random time was selected from a uniform distribution between the 15 and 85 percentiles of the waiting times (time from trial start to EMG onset) of the 100 pedal presses in stage I. We further ensured that there was no EEG evidence for movement preparation at the randomly selected time points by eliciting *RP*− cues only if the classifier output indicated as being within the “Idle” class.

### 2.6. Statistical analysis

We ran two Logistic mixed-effects analyses using the *glmer* function in the lme4 package (Bates, D, Mächler, M, Bolker, B.M., Walker 2015). In both analyses, we used a model comparison approach to select the optimal random effect structure, as suggested in (Matuschek et al. 2017). Exhaustive details of the step-by-step random effect selection process are available in the supplementary material. Below, we provide a summary of the analysis.

#### 2.6.1. Prospective vs. retrospective contributions

The aim of the first analysis was to investigate whether intention reports are influenced by prospective (motor preparation) and retrospective (action execution) components. For this, we used all selected *Go* and *No*-*Go* trials and fit a logistic regression to predict the proportion of awareness reports based on the presence or absence of an RP (*RP*+/*RP*−), the execution or inhibition of an Action (*Go*/*No*-*Go*) and the interaction between both factors (RP × Action).

#### 2.6.2. Dynamic integration of prospective and retrospective cues

The second analysis aimed to study whether retrospective reconstruction and motor preparation interact in a time-dependent manner. For this, we studied selected *Go* trials only, because integration of prospective motor preparation and retrospective sensory feedback was only possible when participants executed an action (i.e. when sensory feedback was present). In *Go* trials, participants could execute an action at different times after cue presentation. We refer to this time delay as reaction time (RT), its characteristics are described in detail in the Results. For each participant, we excluded trials where the RT was above or below 3*SD* from the individual mean. Because we were interested in the effect of absolute time passed between cue presentation and movement initiation rather than the deviation from mean RT, we did not centre the RT predictor variable. Thus, we fit a logistic regression to predict the probability of reporting awareness given the presence or absence of an RP (*RP*+/*RP*−), the continuous reaction time (*RT*) and their interaction (*RP* × *RT*).

### 2.7. EEG-informed selection of participants

The readiness potential is the target brain signal that we used to manipulate prospective information about motor preparation. While the readiness potential is a potentially informative feature used by the classifier, it is not guaranteed *a priori* that the EEG features that were extracted by the classifier in order to separate both classes were based on the presence and absence of an RP over central channels. This, in turn, does not guarantee that *RP*+ and *RP*− cues in stage II were triggered by the presence and absence of RPs, respectively. We thus examined the recorded EEG data of each participant individually and found that for 7 out of the 23 participants this prerequisite was not met. These participants were thus excluded from all analyses. A detailed description of the selection approach is presented in the ***Supplementary Information***.

## 3. Results

In our experiment, participants performed a self-paced movement task and were occasionally interrupted by a cue which instructed them to either execute or inhibit a movement. The timing of cues was based on the motor preparation state determined in real-time by a BCI from the ongoing EEG. Participants were asked to report whether they were intending to move *at the time* the cue appeared. Before we report findings from our main analysis in Section 3.2, in the following we first characterize all recorded trials and explain the criterion by which we selected a subset for the main analysis.

### 3.1. Data description

The number of trials in which participants were presented a cue, as well as the exact times when cues were presented, could not be precisely experimentally controlled. In case of *RP*+ trials, this is because the BCI was calibrated so as to elicit cues preferably during the interval just before a movement, based on the detection of a readiness potential. However, the detection of transient events in the EEG in real-time by means of an asynchronous BCI is only possible with a limited accuracy, bound by the noisy nature of EEG signals. In turn, the timing of cues in *RP*− trials was predetermined at random at trial start. In the following, we characterize all recorded trials by whether and when a movement and/or a cue occurred during each trial, and the trial selection procedure. Fig 2 illustrates the types of trials that occurred during the task and highlights the ones that were included for analysis.

**Figure 2.**
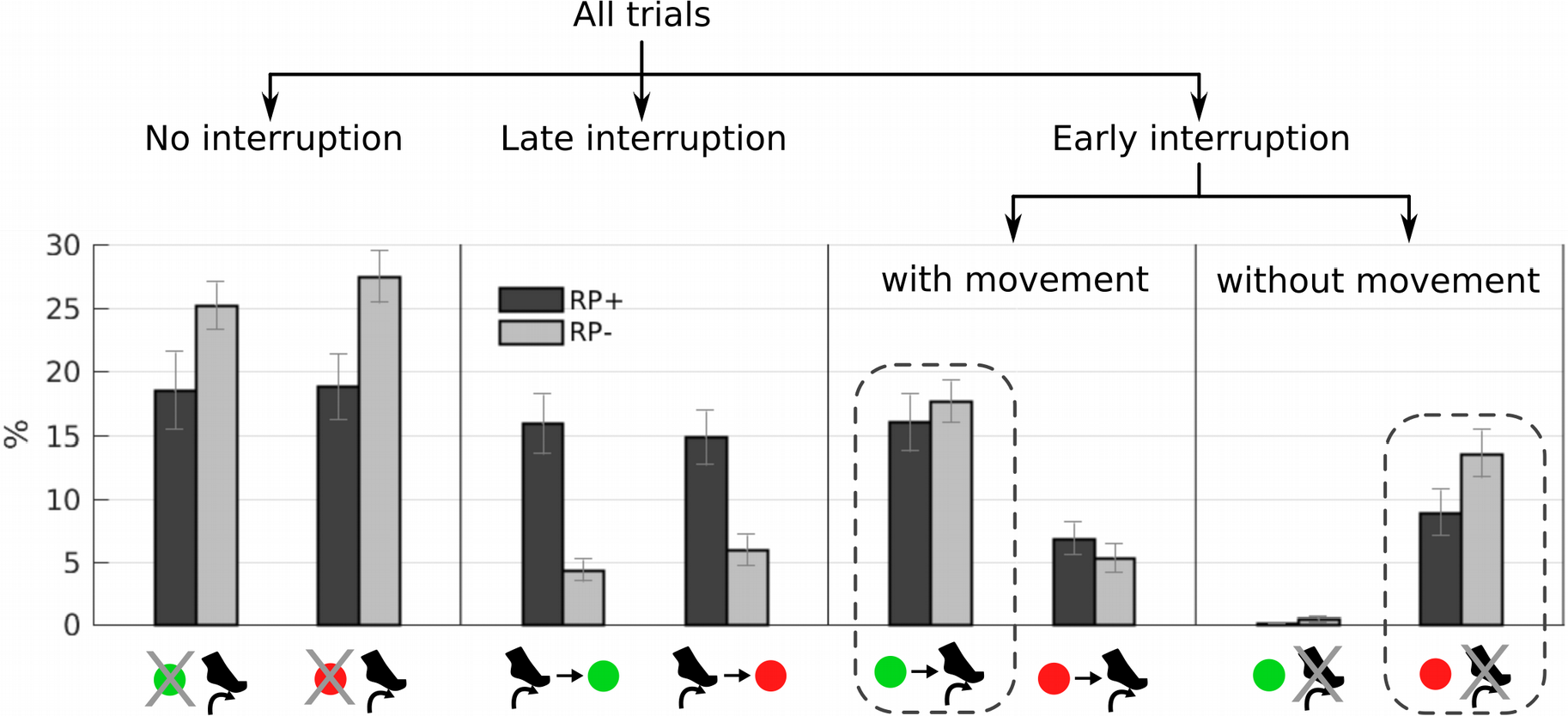
Types of observed trials and selection procedure. Bar graphs represent the grand-averaged percentage (+ SEM) of trials within each category, for Go (green cues) and No-Go (red cues) trials, and in the RP+ (dark grey) and RP− condition (light grey). Percentages are calculated in RP+ and RP− conditions separately, i.e. dark grey and light grey bars sum up to 100, respectively. The pictograms below the bar graphs indicate the temporal relation between cue presentation and movement onset. *No interruption* trials: In some trials, participants executed a movement and no cue was presented at all. In these trials, no awareness report was collected, and no further analysis was conducted. *Late interruption* trials: In some trials, cues came “too late”, shortly after participants had already started moving. All these trials were discarded from further analysis. *Early interruption* trials: Cues were shown before any EMG onset was detected. In the Go condition, only trials where participants moved after the green cue presentation were included for analysis (dashed line). In the No-Go condition, only trials where no EMG onset was detected after the red cue presentation were included for analysis (dashed line).

#### 3.1.1. Characterization of trial types

In some cases, participants pressed the pedal *without* a cue being elicited (*No interruption* trials). In the *RP*+ condition, these represent instances where the BCI failed to detect a readiness potential (*RP+/Go*: *M* = 18.5%, *SEM* = 3.1%; *RP+/No-Go: M* = 18.8%, *SEM* = 2.6%). In *RP*− trials, these represent instances where participants pressed the pedal before the random predetermined time of the cue (*RP−/Go: M* = 25.3%, *SEM* = 1.9%; *RP−/No-Go: M* = 27.5%, *SEM* = 2.0%). In all these cases, since no cue was presented, no awareness report was collected. Thus these trials are excluded from further analysis.

In another subset of trials, a cue was presented *after* EMG onset (*Late interruption* trials). In some *RP*+ trials, a readiness potential was presumably correctly detected by the BCI, but cue was presented after participants had already started moving (*RP+Go: M* = 15.9%, *SEM* = 2.3; *RP+/No-Go: M* = 14.8%, *SEM* = 2.1). In turn, the *RP*− trials where a cue was presented after participants’ movement reflect rare instances where the predetermined probing time by chance coincided with the self-paced time of movement (*RP−/Go*: 4.3%, *SEM* = 0.9; *RP−/No-Go: M* = 5.9%, *SEM* = 1.2). For our purposes, these cues came too late and the corresponding awareness reports are thus excluded from further analysis.

In another subset of trials, the cue was presented *before* EMG onset. In the *Go* condition (*Early interruption* trials *with movement*), these trials fulfil our prerequisite that *Go* cues must be followed by a movement, and thus the corresponding awareness reports are used in the main analysis (*RP+/Go: M* = 16.0%, *SEM* = 2.2; *RP−/Go: M* = 17.7%, *SEM* = 1.7). In the *No*-*Go* condition, participants sometimes *initiated* a movement after a cue was presented (*RP+/No-Go: M* = 6.9%, *SEM* = 1.3; *RP−/No-Go:* 5.3%, *SEM* = 1.1). Although they were often able to abort a movement before fully pressing the pedal in some of these trials, the very initiation of a movement - even though aborted - might suffice for participants to reconstruct an awareness of intention in the awareness probes that followed those cues. Thus, these trials were excluded from further analysis.

Finally, in some trials a cue was elicited before any EMG onset but no movement was produced after it (*Early interruption* trials *without movement*). In the *Go* condition, these very rare occurrences reflect trials where participants failed to respond with a pedal press to a green cue (*RP+/Go*: *M* = 0.15%, *SEM* 0.06; *RP−/Go*: *M* = 0.4%, *SEM* = 0.2). In contrast, as expected, in the *No*-*Go* condition this occurred more frequently (*RP+/ No-Go: M* = 8.9%, *SEM* = 1.8; *RP−/ No-Go: M* = 13.6%, *SEM* = 1.8). In these trials, participants successfully followed the instruction to withhold any movement after a red cue. Because they fulfil our prerequisite that *No*-*Go* cues must not be followed by a movement, the corresponding awareness reports are used in the main analysis.

#### 3.1.2. Time relation between movements and cues

A closer look into the distributions of time differences between EMG onsets and cues provides further insight into the way in which our experimental design resulted in the observed proportions of trials (*Figure 3*).

**Figure 3.**
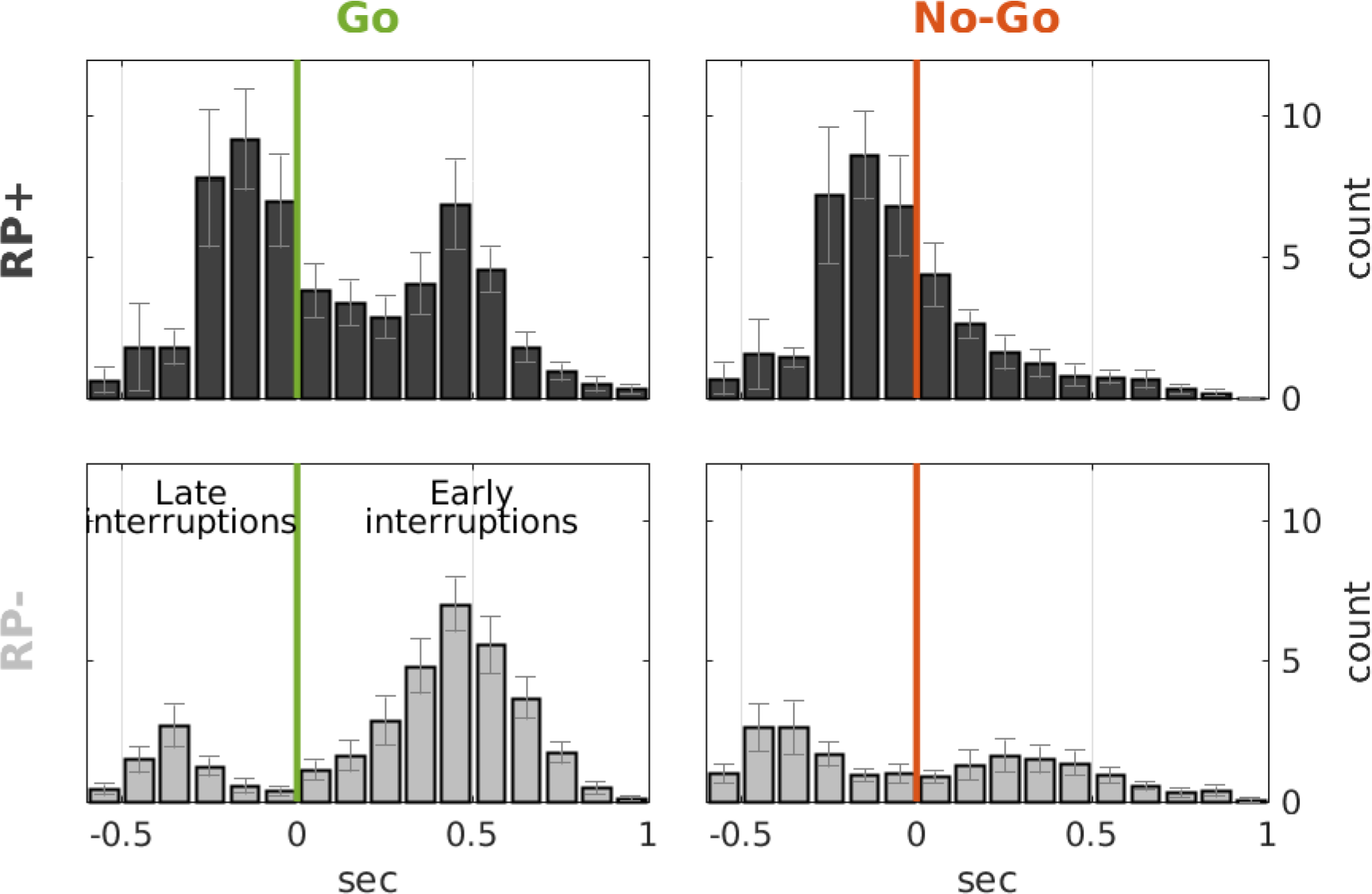
Distribution of EMG onsets with respect to the time of probe presentation in *Go* (left) and *No*-*Go* (right) trials, in the RP+ (top) and RP− condition (bottom). Negative times correspond to the distribution of EMG onsets in *Late interruption* trials, in which the cue was presented after participants started moving. Positive times correspond to the distribution of EMG onsets in *Early interruption* trials (i.e. classic reaction times), where a cue was presented and a movement was initiated shortly afterwards. Bars and antennas show the grand averages and SEMs of trial counts, respectively.

*Go* and *No*-*Go* signals were often triggered *after* participants had started moving in the *RP*+ conditions, while this was rarely the case in *RP*− condition. These *Late interruption* trials correspond to the distribution centered before cue presentation in the RP+ condition. These are instances of motor preparation states that were successfully detected by the BCI, but too late. However, the trials falling on the right tail of this distribution were instead instances where motor preparation was successfully interrupted early by the BCI. These trials can be interpreted as interruptions after the point of no return (Schultze-Kraft et al., 2016). That is, trials in which participants would have moved *anyway* if a cue had not been presented. In fact, in a number of *No*-*Go* trials participants failed to inhibit a movement and an EMG onset was detected after the red cue. In turn, in the *Go* condition, the effect of these intercepted self-paced actions is visible in the higher count of trials with very fast responses (RT < 200 ms) in the *RP+/Go* condition compared to the *RP−/Go* condition. In sum, in the *RP+/Go* condition, very fast trials (<200 ms) include both self-paced movements that happened to occur just after the green *Go* signal (right tail of the “late interruption” distribution), and also reactions to the *Go* signal (left tail of the “early interruptions” distribution; see *Figure 3*). Instead, in the *RP−/Go* condition, movements produced very fast after the cue presentation were only reactions.

#### 3.1.3. Fast RTs in Go trials

We checked that these very fast responses in the left tail of the “early interruption” distribution could physiologically be fast reactions rather than self-paced actions that the classifier did not predict, by looking at the RT distribution to *Go* cues on stage III (*Figure 4*) Reaction times show a skewed distribution that is typical for simple cued reaction time tasks. In that task, where no self-paced actions were being performed and participants were only reacting to *Go* cues presented at random times, we also observed some very fast reaction times (<200 ms) comparable to the ones found in the *RP*− condition in stage II.

**Figure 4.**
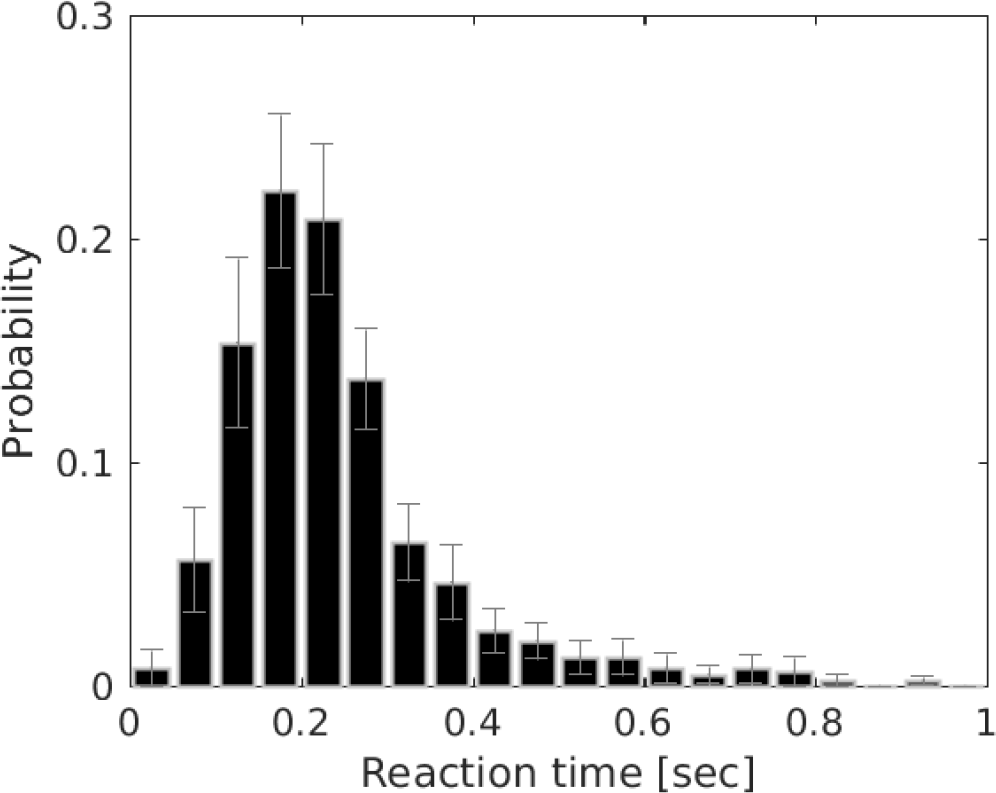
Distribution of reaction times from stage III. The histogram shows, in discrete 50 ms bins, the probability (± SEM) of observing EMG onsets after presentation of the Go cue.

#### 3.1.3. Trial selection for main analysis

As detailed above, the selected subset of trials for the main analysis included only those in which the cue was presented before any movement onset and was either followed by a movement in the *Go* condition or by no EMG onset at all in the *No*-*Go* condition. The analysed trials thus included an average of 29 (*SEM* = 4) *RP*+/*Go* trials, 29 (*SEM* = 3) *RP−/Go* trials, 16 (*SEM* = 3) *RP+/ No-Go* trials and 23 (*SEM* = 3) *RP*−/ *No*-*Go* trials, per participant.

### 3.2. Main analysis

To study how motor preparation processes (RP+/RP−) and action execution processes (Go/No-Go) influence the experience of intention, we fitted two logistic regression models to participants’ responses. In each model we used two independent variables as predictors to test specific predictions made by the retrospective, prospective and temporal integration hypotheses. In each models we used two independent variables as predictors to test specific predictions made by the retrospective, prospective and temporal integration hypotheses (see *Methods* section). the prospective contribution to intention awareness was tested by using the categorical variable motor preparation state (*RP*+/*RP*−) as the first predictor (vertical arrows in Fig 2).

#### 3.2.1. Prospective and retrospective contributions to motor intention awareness

The prospective hypothesis suggests that participants have conscious access to their motor preparatory processes before movement initiation, and it thus predicts affirmative intention judgements to be more likely in the *RP*+ than in the *RP*− condition. In turn, the retrospective hypothesis predicts that the execution of a movement will yield more awareness reports than the absence of an action.

As shown in Figure 5, participants were significantly more likely (*X*^*2*^_*(1)*_ = 21.27, *p* < 0.001) to report awareness in the *Go* (*M* = 35.2%, *SEM* = 6.4%) than in the *No*-*Go* condition (*M* = 16.0%, *SEM* = 4.7%). This suggests a strong effect of retrospection: the presence of an action strongly increased the probability of participant’s reporting an intention to move at the time of probing, compared to trials where no overt movement was present. Furthermore, participants were also significantly more likely (*X*^*2*^_*(1)*_ = 5.04, *p* = 0.024) to report awareness of an intention to move in the *RP*+ (*M* = 32.7%, *SEM* = 6.1%) than in the *RP*− condition (*M* = 24.0%, *SEM* = 6.0%). That is, if they were preparing to move at the time the probe appeared, they were more likely to report an intention than if they were not preparing. No significant interaction was found (*X*^*2*^_*(1)*_ = 1.68, *p* = 0.193).

**Figure 5:**
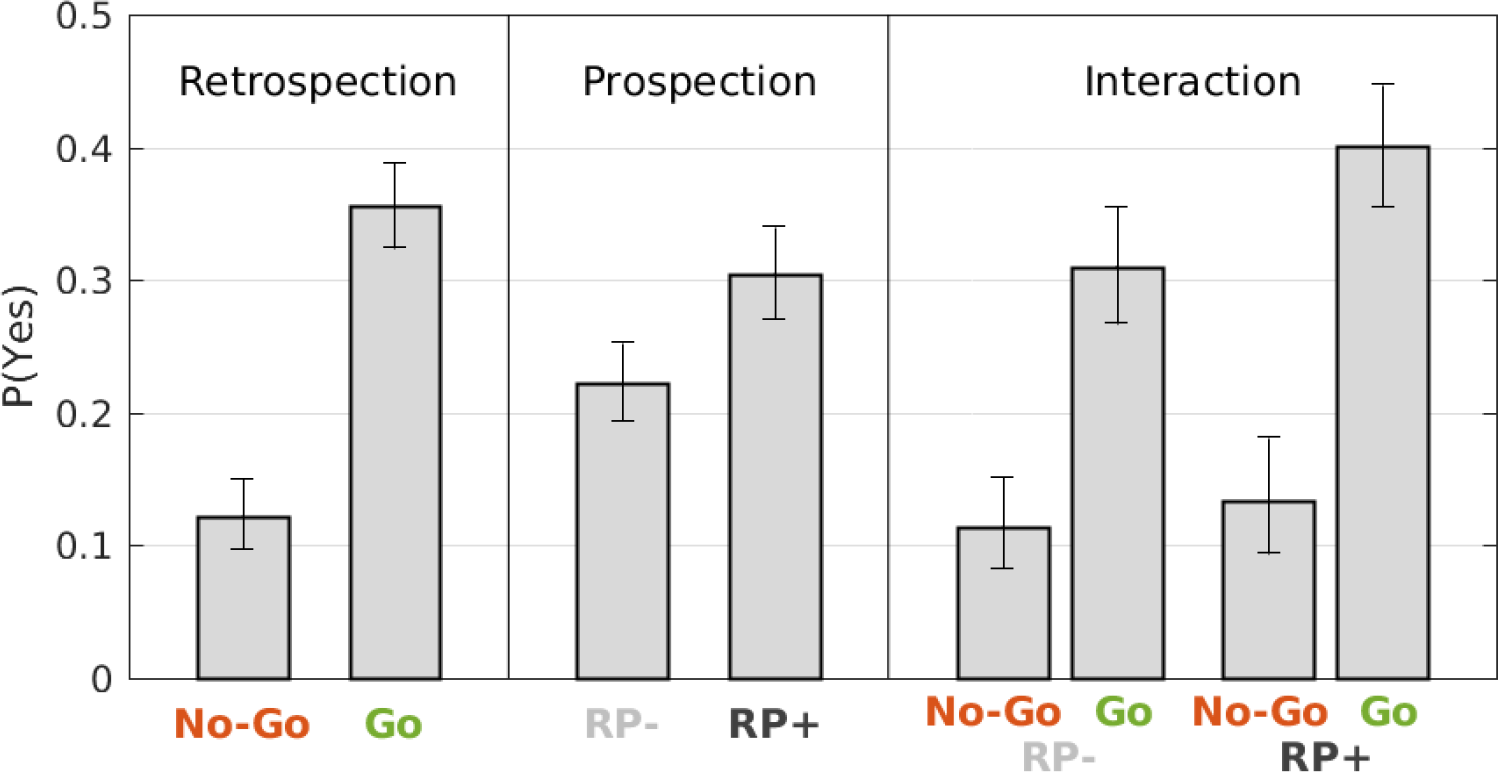
Probability of responding Yes in *No*-*Go* and *Go* trials (left), in *RP*− and *RP*+ trials (middle), and in their corresponding intersections (right). These comparisons reflect the retrospective and prospective contributions (and their interaction) to awareness judgements, respectively. Bars and antennas show probability estimates and 95% confidence intervals, respectively, calculated by pooling the responses of the corresponding subset of trials across all participants.

#### 3.2.2. Dynamic integration of prospective and retrospective information

We next investigated whether prospective and retrospective cues are dynamically integrated in intention awareness judgements. The temporal integration hypothesis suggests that the experience of intention depends on the dynamic integration of motor preparation states and the sensory feedback following action execution. Because *No*-*Go* trials lacked any movement and thus a measureable reaction time, we restricted this analysis to *Go* trials only. First, this allowed us to test whether the RT modulates the retrospective reconstruction of intention. Second, it allowed us to investigate whether the effect of the motor preparation state triggering the cue (*RP*+/*RP*−) on awareness of intention is dependent the execution of the movement, i.e. the RT. We predict that awareness of intention follows a mechanism similar to comparator models of motor control (Wolpert and Kawato 1998). In particular, we predict that prospective information about motor preparation is only available for integration with (retrospective) sensory feedback for a short time. Thus, we expect intention judgments to be modulated by the time delay between the time elapsed between the motor preparation state triggering the cue and the execution of a movement.

As shown in Figure 6, the probability of reporting awareness decreased over time (*X*^*2*^_*(1)* =_ *70.74*, p < 0.001). Participants were very likely to report awareness of intention if they initiated a movement shortly after a cue, but very unlikely to report awareness if they were slow. This was the case both in the *RP*+ and the *RP*− condition. Furthermore, the interaction between the RT and the RP (*X*^*2*^_*(1)* =_ 11.04, *p* < 0.001) indicated that the presence of an RP significantly increased the probability of reporting awareness, but *only if* an action was executed within approximately 250 ms after cue presentation. At a hypothetical RT of 0 s, for example, the probability of reporting awareness predicted by the model is 0.944 in the *RP*+ condition, while it is only 0.788 in the *RP*− condition.

**Figure 6:**
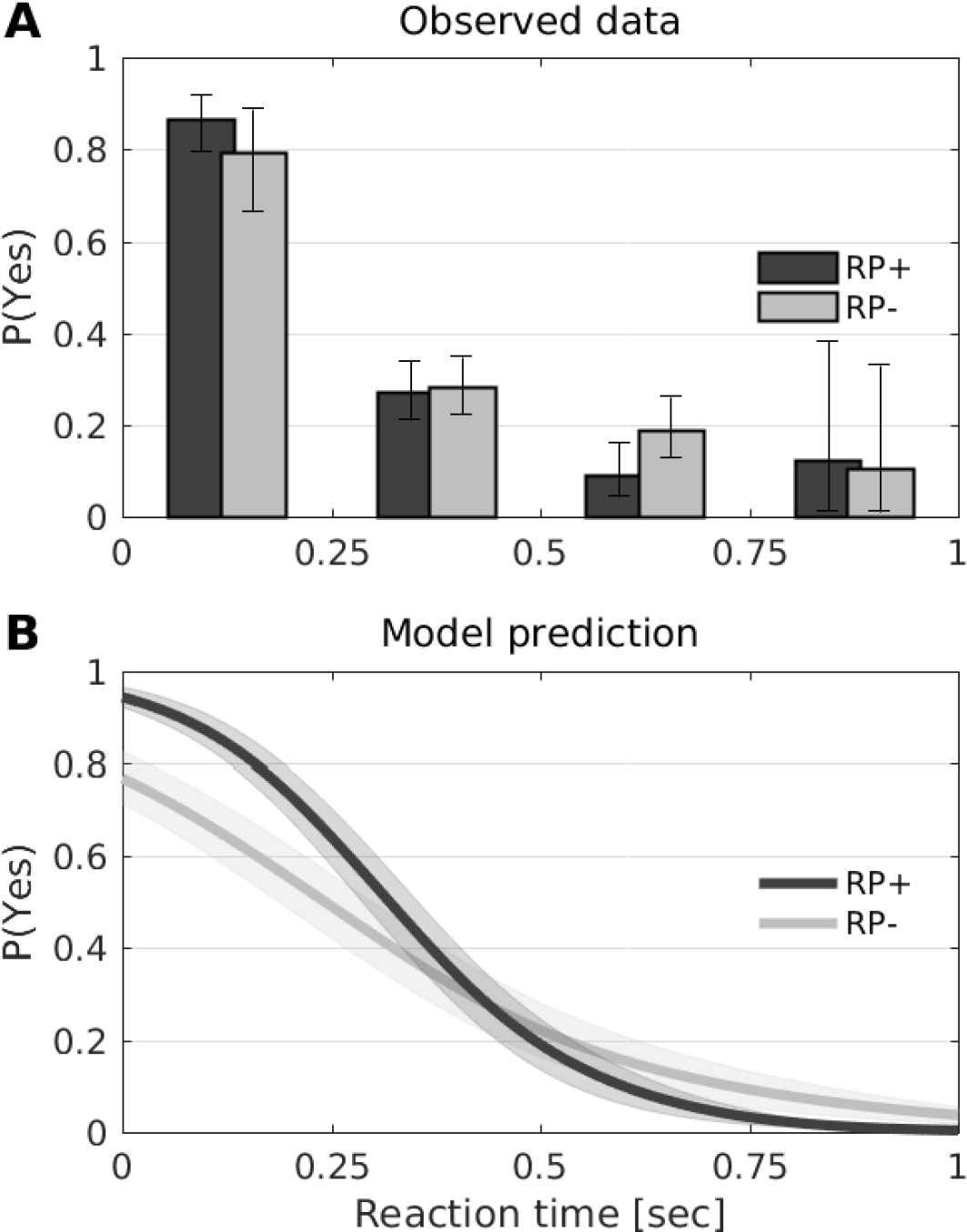
Probability of responding Yes in *Go* trials, for *RP*+ and *RP*− trials individually (color coded) and for different reaction times (x-axis). A. Probability estimates and confidence intervals are calculated by pooling data across participants and are reported according to reaction times in 4 discrete 250 ms bins. B. Model predictions are generated for a continuous reaction time variable. Shown are the grand averages across participants (SEM shown as shaded area).

## 4. Discussion

We conducted an EEG study of intention awareness using a BCI technique which allowed us to monitor motor preparation processes in real-time. Participants performed a self-paced movement task and were occasionally interrupted by a cue which instructed them to either execute or inhibit an action. They were then asked to report whether they were intending to move *at the time* the cue appeared. The time of presentation of the green and red cues was determined by a BCI trained to detect the presence or absence of an RP. This experimental design allowed us to investigate how awareness of intention is manipulated (1) by the presence/absence of motor preparation (prospective hypothesis) (2) by the presence/absence of an action (retrospective hypothesis), and (3) by the potential time lag between the two (temporal integration hypothesis). Our results provide new insight into the elements contributing to the experience of intention.

Our first model showed a strong effect of retrospection. Participants were overall more likely to report an intention to move at the time of probing when an action was executed (*Go* condition) compared to when no overt movement was present (*No*-*Go* condition) (Fig. 5). This is in line with retrospectivist theories suggesting that intention judgements are (at least partially) reconstructed after action execution (Banks and Isham 2009; Kühn and Brass 2009). Further, the estimated probability of reporting awareness was significantly higher for probes preceded by an RP compared to those where no RP was present. This suggests that motor preparation processes influence motor intention perception, both when an action is executed and when it is not. Although the interaction between prospective and retrospective factors was not found to be significant, the data descriptively showed a smaller effect of prospection in the *No*-*Go* condition. Given the limited number of trials available for our analysis, it may be the case that no interaction was detected due to a lack of power rather than due to the absence of an effect.

With our second model, we explored the dynamics of the prospective-retrospective interaction by taking into account the time at which actions were executed with respect to *Go* signals. First, this analysis showed that the retrospective reconstruction of intention found in the first model is time-dependent. Participants were very likely to report awareness when they responded fast to a *Go* signal, but this probability decreased sharply the larger the time difference was between the *Go* signal and movement initiation (Fig. 6). These temporal dynamics were observed in both *RP*+ and *RP*− trials. However, while even in the absence of movement preparation (*RP*−) a mere fast reaction (RT < 250 ms) was sufficient to produce a high percentage of awareness reports (mean=72.8%, SEM=9.5), equally fast trials preceded by movement preparation in the *RP*+ condition resulted in an even higher rate of subsequent intention judgements (mean=86.1%, SEM=5.1). Thus, the model revealed a time-dependent effect of prospection: when a movement was executed, the presence of an RP increased the probability of awareness reports if the movement was executed within approximately 250 ms after a *Go* cue. Because the cues were locked to the presence or absence of an RP, the result can also be phrased as follows: in the *Go* condition the RP only made awareness reports more likely when an action took place shortly after movement preparation was detected. This suggests that information about prospective motor preparation is available for a limited period of time, and that it may be overwritten by retrospective reconstruction. This time-constraint is consistent with everyday experience, and is confirmed by the timings of self-paced actions shown in Fig 2. Normal self-paced actions are typically executed immediately after an RP is present, and not otherwise. Therefore, it makes sense that only events (i.e. actions) happening at a physiologically plausible time after presence of a motor preparation brain signal are integrated with that brain state and perceived as its consequence.

A few considerations are worth noting before the final conclusions. First, this experiment used *delayed* reports of intention awareness. Because we were interested in studying both the prospective and retrospective contributions to motor intention awareness, participants provided their intention judgement after the allowed motor response time window of 1.5 sec. Thus, although we asked participants about their intention at the time of the cue, our results cannot be interpreted as reflecting the experience of intention participants had *at the time* the probe was presented. The RP is a transient signal, and it decays over time. Thus, while the observed effect of the RP on intention awareness observed in the first analysis was small, the possibility remains that the effect of prospection could be higher if reports were obtained closer to the time of action.

Second, awareness of intention in our task was not essential for successful task completion. Participant’s main goal was to respond to the *Go* and *No*-*Go* cues correctly, and intention was only reported afterwards. This contrasts with other awareness report methods, where action execution is contingent on the conscious experience (e.g. Matsuhashi and Hallett 2008; Parés-Pujolràs et al. 2019)) and thus focus on the prospective component of motor awareness. Further research is required to evaluate to what extent purely prospective cues are accessible to guide action using paradigms where task performance is dependent on awareness.

Finally, our results show strong evidence for retrospective reconstruction of intention: participants were more likely to report an intention to move at the time of a cue if they executed a movement following such cue. This effect is time-dependent: they are more likely to report an intention if they respond fast to the cue, and unlikely to report it if their reaction to the cue is slow. This is consistent with retrospectivist theories suggesting that judgements of intention are (at least partially) reconstructed after action execution (Wegner, 2002; Banks and Isham 2009; Kühn and Brass 2009). However, one question remains open. Assuming that participants have a certain “baseline” probability of reporting awareness (i.e. based on prospective cues or individual biases), two mechanisms might explain the observed retrospection results. First, it could be the case that execution of an action *increases* the probability of reporting an intention, as the aforementioned theories suggest. This would be a type of “positive reconstruction”, where people “make up” intentions based on external events. However, it is also possible that, instead, inhibition of an action *decreases* the probability of reporting awareness. That is, there could be a kind of “negative reconstruction” in which participant’s intention reports become less likely (i.e. due to loss of sensitivity to prospective cues, or increased reliance on external cues – or the lack thereof). Both mechanisms might coexist. While the present data show that some kind of retrospective effect is present, it cannot disentangle these two possibilities because no “baseline” awareness rate is available. Future research might attempt to investigate this question by developing a method that could provide a “baseline” estimate of the likelihood of reporting awareness (e.g. verbal reports) and comparing them, within subjects, to reports provided after action execution or action inhibition.

In sum, in this study, we show that both prospective and retrospective cues influence delayed motor intention judgements, and that their integration is time-dependent. The presence of a motor preparation signal increases the probability of reporting an intention (prospective effect) and reports of intention are more likely after executing an action than in the absence of overt movement (retrospective effect). Further, the retrospective effect is modulated by response time (i.e. the probability of reporting awareness decreases the slower the RT), and there is a critical time window of approximately 250 ms during which prospective information is integrated with action execution feedback and influences awareness reports. This research sheds new light into the prospective and retrospective components of awareness of intention, and provides new methods to investigate the neural correlates of voluntary action execution and the related experiences of intention and agency.

## 5. Supplementary Information

### 5.1. EEG-informed selection of participants

The readiness potential was the target brain signal that we aimed to use in order to manipulate prospective information about motor preparation. Therefore, we investigated whether such BCI-based manipulation was effective in each individual participant.

A qualitative assessment of EEG data from stage I used to train the classifier (Figure S1A) shows that for most participants the signals look as expected, with EEG signals preceding self-paced movements in average displaying the typical negative trend of a readiness potential (“Move” class), while EEG signals preceding trial start cues do not show any particular trend (“Idle” class). However, while the RP is a potentially informative feature that the BCI may use for classification, it is not guaranteed a priori that the EEG features extracted by the classifier to separate the “Move” and “Idle” classes were based on the presence and absence of an RP over central channels.

A visual inspection of stage II data (Figure S1B) suggests that for most participants *RP*+ cues were effectively preceded by an RP-like negativity, while *RP*− cues were not (with some conspicuous exceptions). Note that here we only consider *RP*+ and *RP*− cues that were elicited *before* any movement onset, thus excluding EEG data that would otherwise be contaminated with signals related to movement execution. In order to test whether we could rely on the BCI-triggered cues during stage II to discriminate *RP*+ from *RP*− activity in each individual participant, we performed the following analysis. Channel Cz was chosen for analysis because readiness potentials preceding foot movements are typically most distinct over that channel (Brunia et al., 1985, Schultze-Kraft et al., 2016). For each trial individually, we subtracted the average EEG signal in the time interval −200ms to 0ms from the average EEG signal in the time interval −1200ms to −1000ms, with respect to the time of cue presentation. These values represent the relative change in amplitude in channel Cz during the 1.2 seconds before cue presentation. If the BCI relied on the readiness potential for classification, EEG signals over central channels preceding *RP*+ cues should be on average more negative than signals preceding *RP*− cues. To test this hypothesis, we ran a two-sample one-sided t-test, for each participant separately. The box plots in Fig S1C show, for each participant and for *RP*+ and *RP*− cues individually, the distributions of amplitude changes, with participants ordered by the t-statistics of the t-test from largest (left) to smallest (right). For the first 16 participants the t-test showed that signals preceding *RP*+ cues became significantly more negative during the 1.2 sec interval than signals preceding *RP*− cues. For the remaining 7 participants this was not the case, suggesting that the classifier did not made predictions based on the presence or absence of RP-like events in the EEG. Consequently, these participants are excluded from all subsequent analyses. Fig S1D shows individual and grand average EEG signals preceding *RP*+ and *RP*− cues of the 16 participants selected for the final sample.

**Figure S1:**
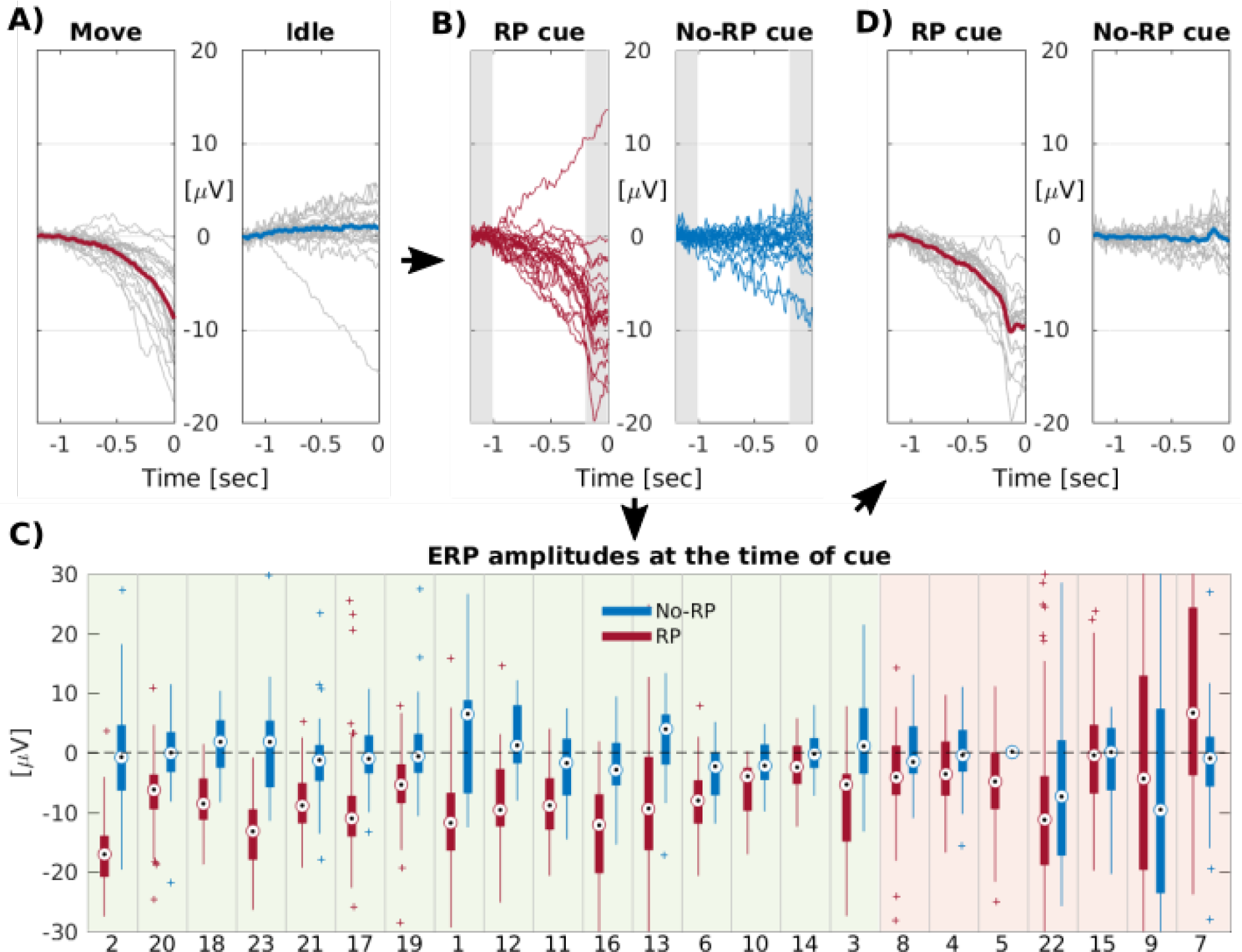
Selection of suitable participants based on EEG signals of channel Cz. **A.** Event-related potentials (ERPs) of EEG signals recorded during stage I, time-locked to self-paced EMG onsets (left, “Move”) and time-locked to trial start cues (right, “Idle”). ERPs are baseline corrected in the interval −1200 to −1000 ms, and shown for individual participants (grey) and as grand average (colored). **B.** ERPs of EEG signals recorded in stage II, time-locked to *RP*+ cues (left) and time-locked to *RP-* cues (right). ERPs are baseline corrected in the interval −1200 to −1000 ms and shown for individual participants. **C.** For each participant (ID on x-axis) and for *RP*+ and *RP*− cues individually (color coded), the box plots show the distribution of EEG signal amplitude changes between the time interval −1200 to −1000 ms and −200 to 0 ms with respect to cue onset (indicated by gray areas in panel B). Participants are ordered in ascending order by the t-statistic of a two-sample one-sided t-test that tests whether the mean change in *RP*+ trials was more negative than in *RP*− trials. Participants for which p<.01 are highlighted in green, otherwise in red. **D.** As in B, but only for the selected N=16 participants (gray) and the corresponding grand average (colored).

### 5.2. Model selection procedure and statistical details

As described in the Statistical Analysis section of the methods, we used linear mixed-effects models to test the effects of our explanatory variables on the probability of participants reporting awareness. To select the model that best explained our observed results, we followed the random effect selection procedure suggested in (Matuschek et al. 2017).

In all models, a random intercept was included to account for the variability in the dependent variable across participants. Further, we included those random effects that significantly improved the model fit. To determine the optimal random effects structure, we fit a baseline model which included all explanatory variables and all possible interactions as fixed terms. We then iteratively compared this baseline models against models with one additional random slope using a chi-squared test. If the inclusion of a random slope significantly improved the model fit, the random slope was included in the final model. This approach has been suggested as a better option than including random slopes for all fixed effects, as it decreases the probability of Type II errors while maintaining the same power against type I errors, and has previously been used in the literature (e.g. Steinemann et al. 2018). All models were fit using restricted maximum likelihood estimation (REML) with the *glmer* function in the homonymous R package (Bates, D, Mächler, M, Bolker, B.M., Walker 2015).

Tables S1 and S2 provide the detailed results of the random effect selection procedure for both main analyses and the final inference statistics reported in the main text.

**Table S1.** Model 1 selection steps and statistical results of model comparison. We determined the optimal random effects structure with REML estimation. Random slopes for both RP and Action significantly improved the fit of the baseline model and were therefore included in the model.

| Supplementary Table 1: model 1 random effects selection |  |  |  |
| --- | --- | --- | --- |
| Test individual random effects | Baseline model: |  |  |
|  | P(yes) ~ 1 + RP + Action + RP:Action + (1+\|sub) |  |  |
|  | X<sup>2</sup> | DF | p-value |
| yes ~ 1 + RP + Action + RP:Action + (1+RP\|sub) | 11.79 | 2 | 0.0027 ** |
| yes ~ 1 + RP + Action + RP:Action + (1+Action\|sub) | 17.18 | 2 | <0.001 *** |

**Table S2.**
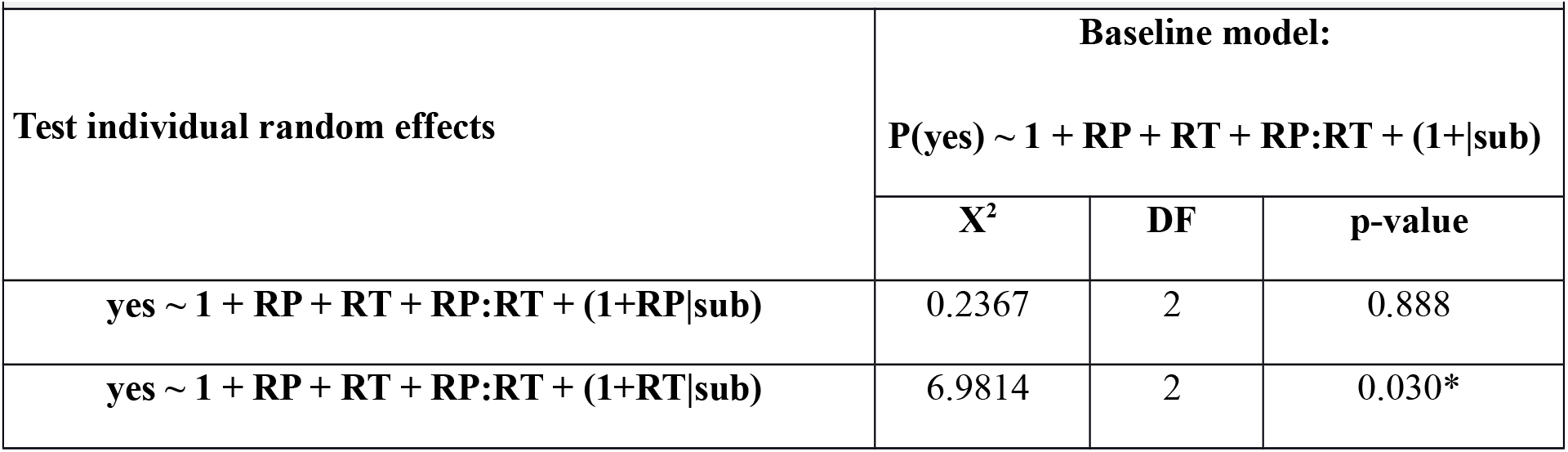
Model 2 random effect selection steps and statistical results of model comparison. We determined the optimal random effects structure with REML estimation. Only the RT random slope significantly improved the fit of the baseline model and was therefore the only random effect included in the model.

| Supplementary Table 2: model 2 random effects selection |  |  |  |
| --- | --- | --- | --- |
| Test individual random effects | Baseline model: |  |  |
|  | P(yes) ~ 1 + RP + RT + RP:RT + (1+\|sub) |  |  |
|  | X<sup>2</sup> | DF | p-value |
| yes ~ 1 + RP + RT + RP:RT + (1+RP\|sub) | 0.2367 | 2 | 0.888 |
| yes ~ 1 + RP + RT + RP:RT + (1+RT\|sub) | 6.9814 | 2 | 0.030* |

## 6. Acknowledgments

This work was partly supported by a joint grant from the John Templeton Foundation and the Fetzer Institute. The opinions expressed in this publication are those of the author(s) and do not necessarily reflect the views of the John Templeton Foundation or the Fetzer Institute. Furthermore, this work was supported by the DFG Grant Excellence Cluster “Science of Intelligence” and the DFG Grant SFB 940 Volition and Cognitive Control.

